## Supplementary Information for "A Novel Membrane Protein in the *Rhodobacter sphaeroides* LH1-RC Photocomplex"

### Materials and Methods

**Preparation and Characterization of the Native LH1-RC Complex.** The *Rba. sphaeroides* f. sp. *denitrificans* (strain IL106) cells were cultivated phototrophically (anoxic/light) at room temperature for 7 days under incandescent light (60W). This strain can also grow chemoheterotrophically using nitrate and dimethyl sulfoxide (DMSO) as the terminal electron acceptors (dark/anoxic)[1]. Preparation of the native LH1-RC was conducted by solubilizing chromatophores ($OD_{870\text{-nm}}$ = 40) with 1.0 % w/v *n*-dodecyl-β-D-maltopyranoside (DDM) in 20 mM Tris-HCl (pH 8.0) buffer for 60 min at room temperature, followed by differential centrifugation. The supernatant was loaded onto a DEAE column (Toyopearl 650S, TOSOH) equilibrated at 4 ˚C with 20 mM Tris-HCl buffer (pH 8.0) containing 0.1 % w/v DDM. The fractions were eluted in an order of LH2, monomeric LH1-RC and dimeric LH1-RC by a linear gradient of NaCl from 0 mM to 400 mM. The peak fractions of monomeric LH1-RC were collected and further purified by sucrose gradient density centrifugation with five-stepwise sucrose concentrations (10, 17.5, 25, 32.5 and 40% w/v) in 20 mM Tris-HCl buffer (pH 8.0) containing 0.05 % w/v DDM at 4 °C and 150,000×g for 6 hours. The monomeric LH1-RC fractions were concentrated for absorption measurement and assessed by negative-stain EM using a JEM-1011 instrument (JEOL) (Supplementary Fig. 1a). Masses and composition of the LH1 and PufX polypeptides were measured by matrix-assisted laser desorption/ionization time-of-flight mass spectroscopy

(MALDI-TOF/MS) and reverse-phase HPLC (Supplementary Fig. 6b, 6c), respectively, using the methods described elsewhere[2]. Quinones were extracted from chromatophores, purified LH1-RC and RC-only complexes, and quinone contents were analyzed (Supplementary Fig. 6d) by the methods described previously[3]. Approximately six UQ-10 molecules were estimated per purified LH1-RC.

**Mutagenesis and Characterization.** A 0.8-kb DNA region at the direct upstream and a 0.9-kb DNA region at the direct downstream of the protein-U gene were amplified by PCR using *Rba. sphaeroides* IL106 genomic DNA as the template. These DNA fragments were connected in a manner that exclude the protein-U gene and cloned in a suicide vector pJPCm[4] using an In-Fusion HD Cloning Kit (TAKARA BIO, Shiga, Japan) as shown in Supplementary Fig. 7. This plasmid was named pJPCm_ΔU, and then a DNA fragment containing *sacRB* genes (lethal genes under the presence of sucrose) and a kanamycin-resistant cartridge was inserted at the unique *Sac*I restriction site on this plasmid. This plasmid was named pJPCm_ΔU-SKm and introduced from the *E. coli* S17-1 λpir host cells by conjugational transfer into the cells of *Rba. sphaeroides* IL106 wild-type strain with a spontaneous resistance to rifampicin. The gene encoding protein-U was deleted from the *Rba. sphaeroides* genome *via* a two-step homologous recombination, which was screened by kanamycin resistance in the first step and by sucrose resistance in the second, as previously described[4]. After the conjugation *E. coli* cells were eliminated by addition of 50 μg/ ml of rifampicin to the growth medium. The removal of the protein-U gene without any insertions of antibiotics-resistant cartridges was confirmed by PCR and DNA sequencing experiments. This mutant was named IL106-ΔU.

Cells of *Rba. sphaeroides* IL106-ΔU were cultivated phototrophically at room temperature under the same condition as that for the wild-type. The chromatophores of both wild-type and IL106-ΔU were treated with 1.0 % w/v DDM in 20 mM Tris-HCl (pH 8.0) buffer for 60 min at room temperature, followed by differential centrifugation. The supernatants (1 mL) with adjusted concentrations of $OD_{870\text{-nm}}$ = 5, 9, 14 were loaded on a stepwise sucrose gradient (10, 17.5, 25, 32.5 and 40% w/v) in 20 mM Tris-HCl buffer (pH 8.0) containing 0.05 % w/v DDM. Centrifugation was conducted at 4 °C and 150,000×g for 6 hours. All of the dimeric and monomeric LH1-RC layers were carefully collected and a ratio of the dimeric to monomeric LH1-RC was calculated using their volumes and absorption intensities (Supplementary Fig. 8).

**Cryo-EM Data Collection.** Proteins for cryo-EM were concentrated to ~3 mg/ml. Two microliters of the protein solution were applied on a glow-discharged holey carbon grids (200 mesh Quantifoil R2/2 molybdenum), which had been treated with $H_2$ and $O_2$ mixtures in a Solarus plasma cleaner (Gatan, Pleasanton, USA) for 30 s and then blotted, and plunged into liquid ethane at –182 ˚C using an EM GP2 plunger (Leica, Microsystems, Vienna, Austria). The applied parameters were a blotting time of 6 s at 80% humidity and 4°C. Data were collected on a Talos Arctica (Thermo Fisher Scientific, Hillsboro, USA) electron microscope at 200 kV equipped with a Falcon 3 camera (Thermo Fisher Scientific) (Supplementary Fig. 1). Movies were recorded using EPU software (Thermo Fisher Scientific) at a nominal magnification of 92 k in counting mode and a pixel size of 1.094 Å at the specimen level with a dose rate of 0.98 e- per physical pixel per second, corresponding to 0.82 e- per $Å^2$ per second at the specimen level. The exposure time was 51 s, resulting in an accumulated dose of 42 e- per $Å^2$. Each movie includes 40 fractioned frames.

**Image Processing.** All of the stacked frames were subjected to motion correction with MotionCor2[5]. Defocus was estimated using CTFFIND4[6]. A total of 551,846 particles were selected from 2,766 micrographs using the EMAN2 suite (Supplementary Fig. 2)[7]. The initial 3-D model was generated with 36,284 particles from 87 selected micrographs with underfocus values ranging between 2 and 3 μm using EMAN2. All of the picked particles were further analyzed with RELION3.0 and 3.1[8], and 219,927 particles were selected by 2-D classification and divided into four classes by 3-D classification resulting in only one good class containing 160,488 particles. The 3-D auto refinement without any imposed symmetry (C1) produced a map at 3.10 Å resolution after contrast transfer function refinement, Bayesian polishing, masking, and post-processing. The selected 160,448 particle projections were subjected to subtraction of the detergent micelle density followed by 3-D auto refinement to yield the final map with a resolution of 2.94 Å according to the gold-standard Fourier shell correlation using a criterion of 0.143 (Supplementary Fig. 2)[9]. The local resolution maps were calculated on RESMAP[10].

**Model Building and Refinement of the LH1-RC Complex.** The atomic models of the LH1 ring of the *Tch. tepidum* LH1-RC (PDB: 5Y5S) and the RC of the *Rba. sphaeroides* (PDB: 1PCR) was fitted to the cryo-EM map obtained for the *Rba. sphaeroides* LH1-RC using Chimera[11]. Amino acid substitutions and real space refinement for the peptides and cofactors were performed using COOT[12]. Whole regions of PufX and protein-U as well as both terminal regions of the LH1 α-subunit were modelled *ab-initio* based on the density. The manually modified model was refined in real-space on PHENIX[13], and the COOT/PHENIX refinement was iterated until the refinements converged. Finally, the statistics calculated using MolProbity[14] were checked. Figures were drawn with the Pymol Molecular Graphic System (Schrödinger)[15] and UCSF Chimera[11].

**Supplementary Table 1 Cryo-EM data collection, refinement and validation statistics.**

| | LH1-RC-PufX-Protein-U complex (EMDB-31400, PDB ID: 7F0L) |
|---|---|
| **Data collection and processing** | |
| Magnification | 92000 |
| Voltage (kV) | 200 |
| Electron exposure (e–/Å$^2$) | 42 |
| Defocus range (μm) | -0.7 to -2.6 |
| Pixel size (Å) | 1.094 |
| Symmetry imposed | C1 |
| Initial particle images (no.) | 551846 |
| Final particle images (no.) | 160488 |
| Map resolution (Å) | 2.9 |
| FSC threshold | 0.143 |
| Map resolution range (Å) | 313-2.9 |
| **Refinement** | |
| Initial model used (PDB code) | 5Y5S, 1PCR |
| Model resolution (Å) | 3.1 |
| FSC threshold | 0.5 |
| Model resolution range (Å) | 140-2.9 |
| Map sharpening *B* factor (Å$^2$) | -63 |
| Model composition | |
| Non-hydrogen atoms | 23458 |
| Protein residues | 2277 |
| Ligands | 112 |
| *B* factors (Å$^2$) | |
| Protein | 31.4 |
| Ligand | 33.3 |
| R.m.s. deviations | |
| Bond lengths (Å) | 0.007 |
| Bond angles (°) | 2.731 |
| Validation | |
| MolProbity score | 1.93 |
| Clashscore | 10.74 |
| Poor rotamers (%) | 1.93 |
| Ramachandran plot | |
| Favored (%) | 97.09 |
| Allowed (%) | 2.91 |
| Disallowed (%) | 0.00 |

**Supplementary Table 2 Comparison of the distances of His–BChl(Mg) and BChl(Mg)–BChl(Mg) in LH1, LH2 and RC special pairs from various phototrophic bacteria.**

| | Distance of His(Nε2) to BChl–Mg (Å)[a] | | Distance of Mg–Mg (Å)[a] | |
|---|---|---|---|---|
| LH1 or LH2 | α | β | Long | Short |
| ***Rba. sphaeroides* (LH1)** | **2.58** | **2.20** | **9.55** | **8.37** |
| *Rsp. rubrum* (LH1) | 2.27 | 2.03 | 9.34 | 8.51 |
| *Rps. palustris* (LH1-W) | 2.93 | 2.71 | 9.61 | 8.29 |
| *Tch. tepidum* (LH1) | 2.19 | 2.19 | 8.88 | 8.72 |
| *Trv.* strain 970 (LH1) | 2.33 | 2.31 | 8.90 | 8.46 |
| *Blc. viridis* (LH1) | 2.54 | 2.25 | 8.8 | 8.5 |
| *Rfx. castenholzii* (B880) | 2.32 | 2.29 | 9.5 | 9.3 |
| *Rps. acidophila* (B850) | 2.34 | 2.34 | 9.5 | 8.8 |
| *Phs. molischianum* (B850) | 2.27 | 2.32 | 9.2 | 8.9 |

| RC (special pair) | L-subunit | M-subunit | BChl *a* (L)–BChl *a* (M) |
|---|---|---|---|
| ***Rba. sphaeroides*** (LH1-RC) | **2.21** | **2.11** | **7.79** |
| *Rba. sphaeroides* (RC-only) | 2.27 | 2.06 | 7.84 |
| *Rsp. rubrum* | 2.09 | 2.12 | 7.76 |
| *Rps. palustris* | 2.73 | 2.74 | 7.69 |
| *Tch. tepidum* | 2.17 | 2.19 | 7.87 |
| *Trv.* strain 970 | 2.33 | 2.31 | 7.65 |
| *Blc. viridis* | 2.36 | 2.35 | 7.83 |

These values were derived from Protein Data Bank: 5Y5S for *Tch. tepidum*, 7C9R for *Trv.* strain 970, 6Z5S for *Rps. palustris*, 6ET5 for *Blc. viridis*, 5YQ7 for *Rfx. castenholzii*, 7EQD for *Rsp. rubrum*, 1NKZ for *Rps. acidophila*, 1LGH for *Phs. molischianum*, 2J8C for *Rba. sphaeroides* (RC-only).

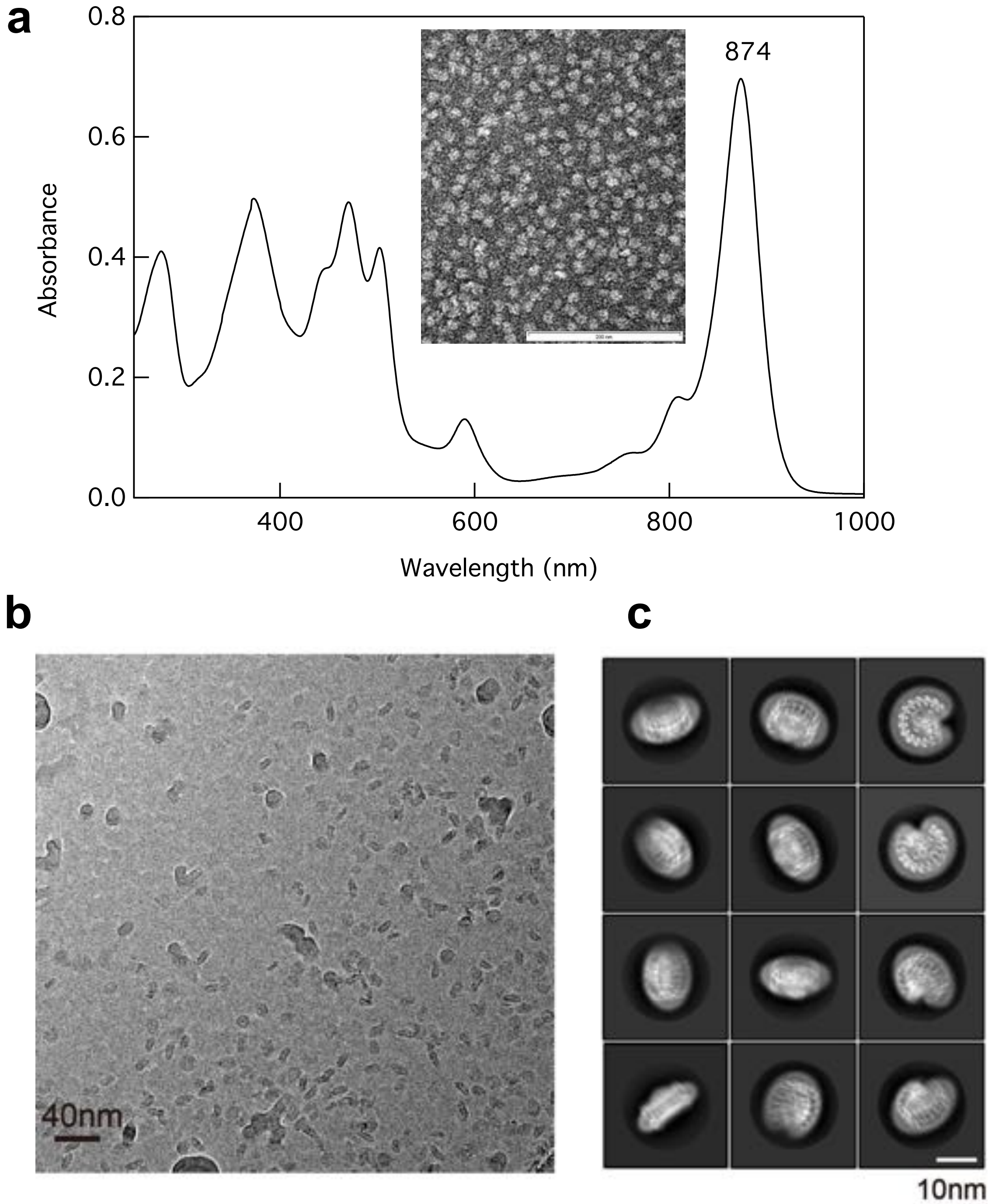

**Supplementary Fig. 1 Absorption spectrum and cryo-EM of the *Rba. sphaeroides* IL106 monomeric LH1-RC complex.** (**a**) Absorption spectrum of the purified monomeric LH1-RC a room temperature. Inset shows negatively stained LH1-RC particles obtained with 0.04 mg/mL LH1-RC in 20mM Tris-HCl (pH7.5) containing .05% DDM. Scale bar: 200 nm. (**b**) A representative cryo-EM micrograph. (**c**) Representative 2D class averages processed from the micrographs of LH1-RC.

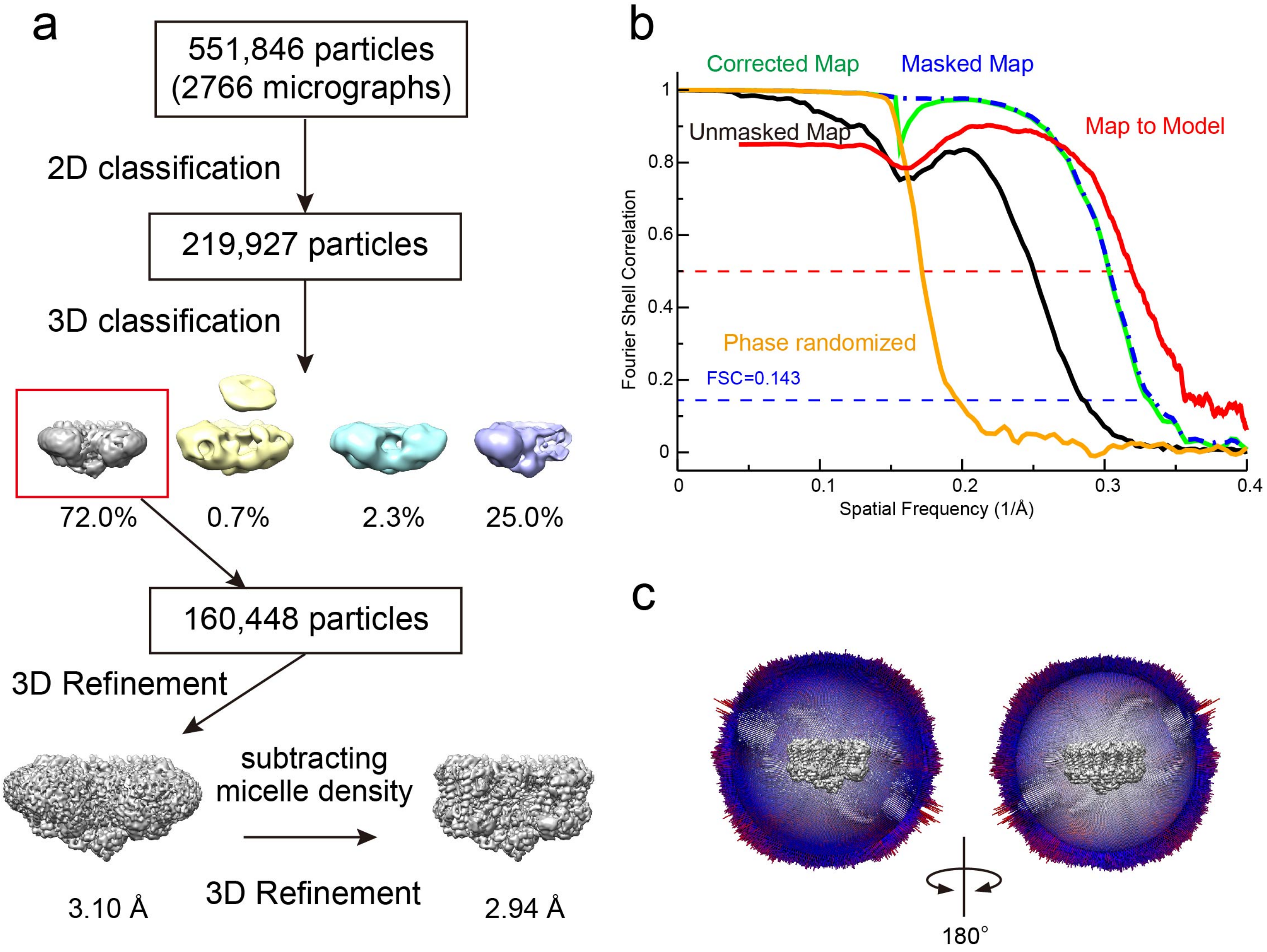

**Supplementary Fig. 2 Structure determination of the *Rba. sphaeroides* IL106 monomeric LH1-RC complex. (a)** Image processing flow of 3D classification and reconstruction. **(b)** The Fourier shell correlation (FSC) plot of the cryo-EM map (unmasked: black, masked: blue, phase randomized corrected: green, phase randomized: orange) and the FSC plot of the model versus the final map (red) are superimposed. **(c)** Angular distribution of reconstructed particles in the C1 map of LH1-RC complex. For clarity, the front half of angular distribution is removed.

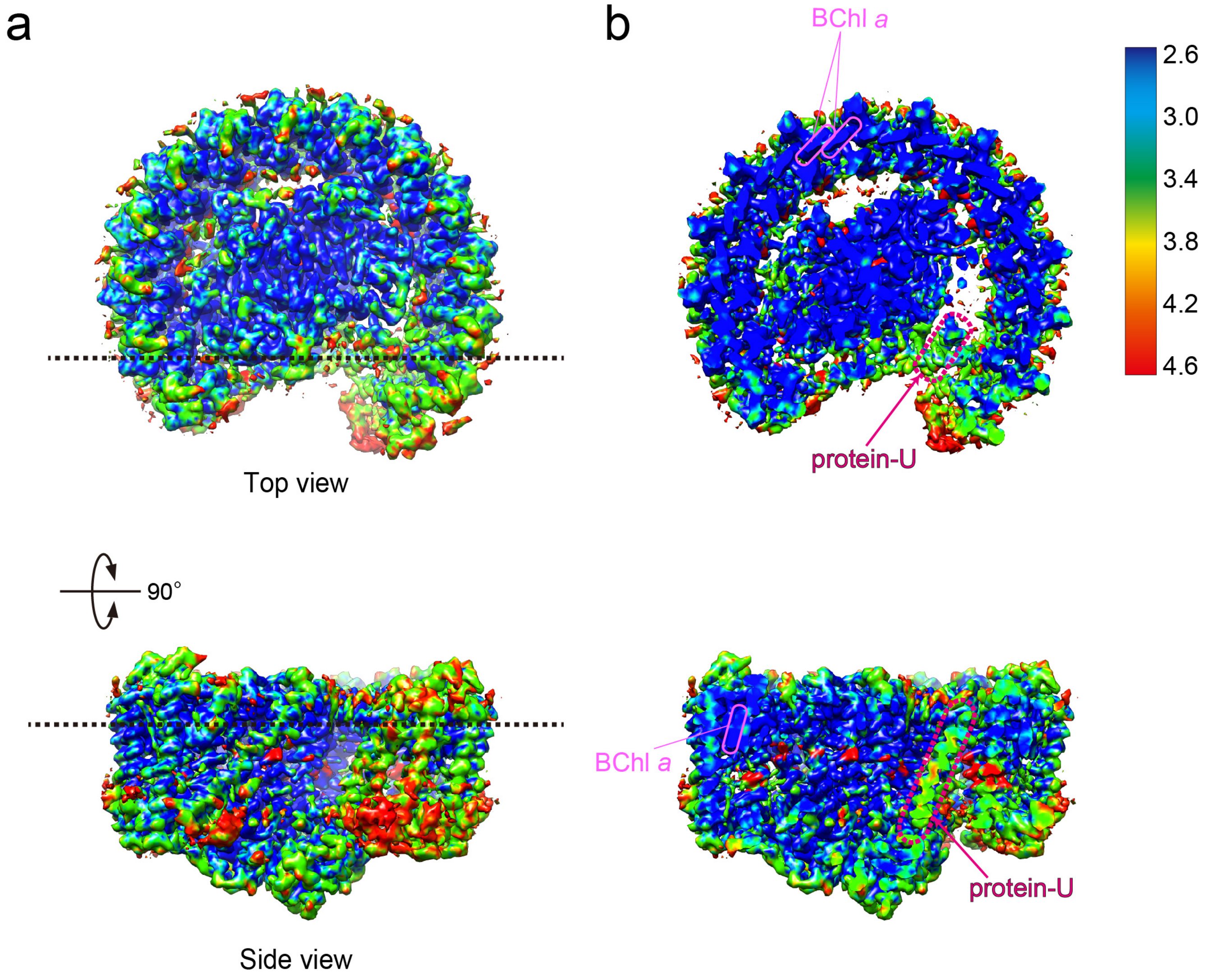

**Supplementary Fig. 3 Local resolution representation of the structure of LH1-RC complex. (a)** Top view from periplasmic side and side view parallel to the membrane plane. Each dotted line indicates the cross section line. **(b)** A central cross sectional view of the left side of the panel. A region indicated by circular pink line and magenta dotted line corresponds to BChl *a* and protein-U, respectively. The map is shown in the colors of the rainbow according to the estimated resolution from 4.6 Å (red) to 2.6 Å (blue).

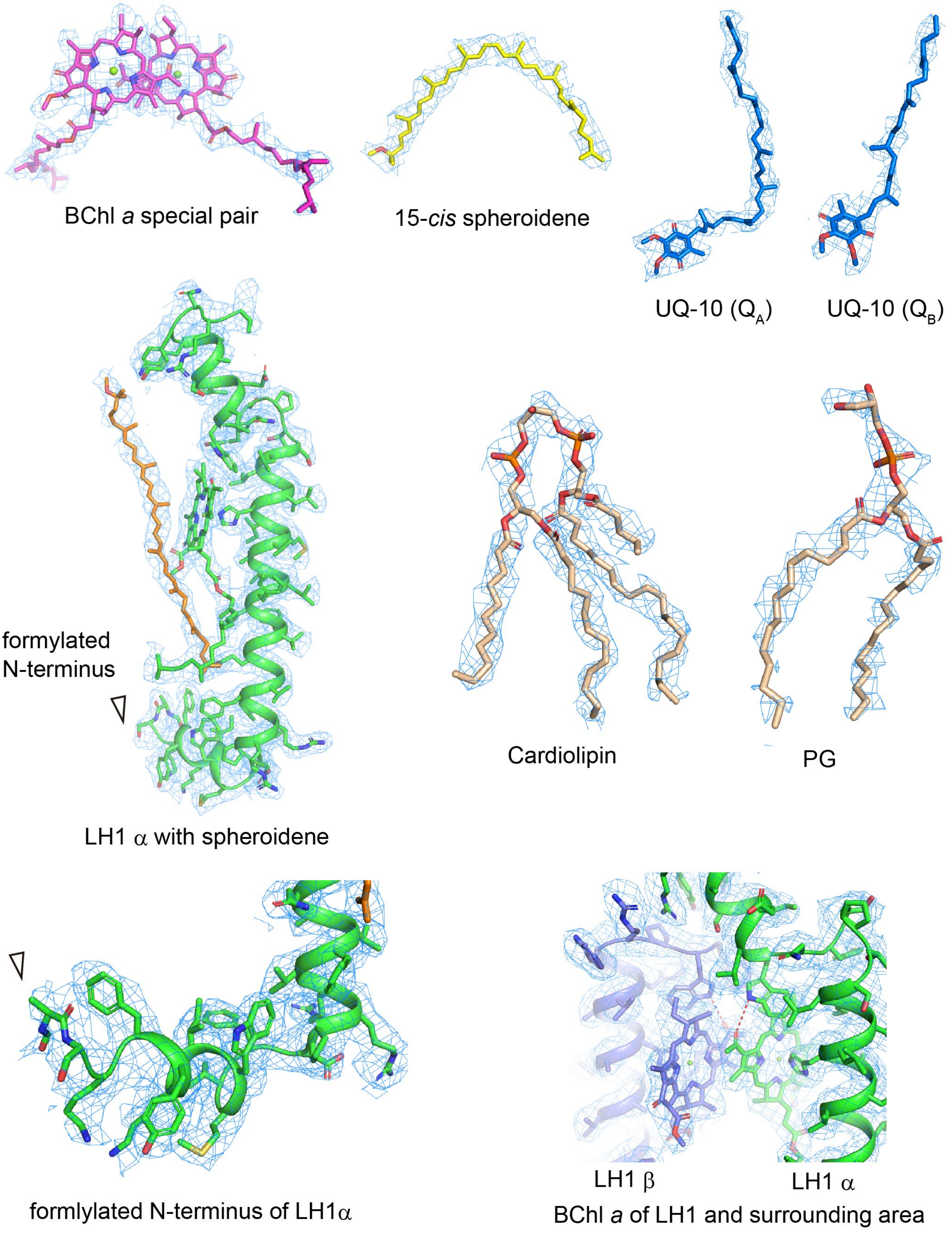

**Supplementary Fig. 4 Cryo-EM densities and structural models in the the *Rba. sphaeroides* IL106 LH1-RC complex.** The color codes of polypeptides are the same as in Fig. 1. The density maps are shown at a contour level of 3.0σ.

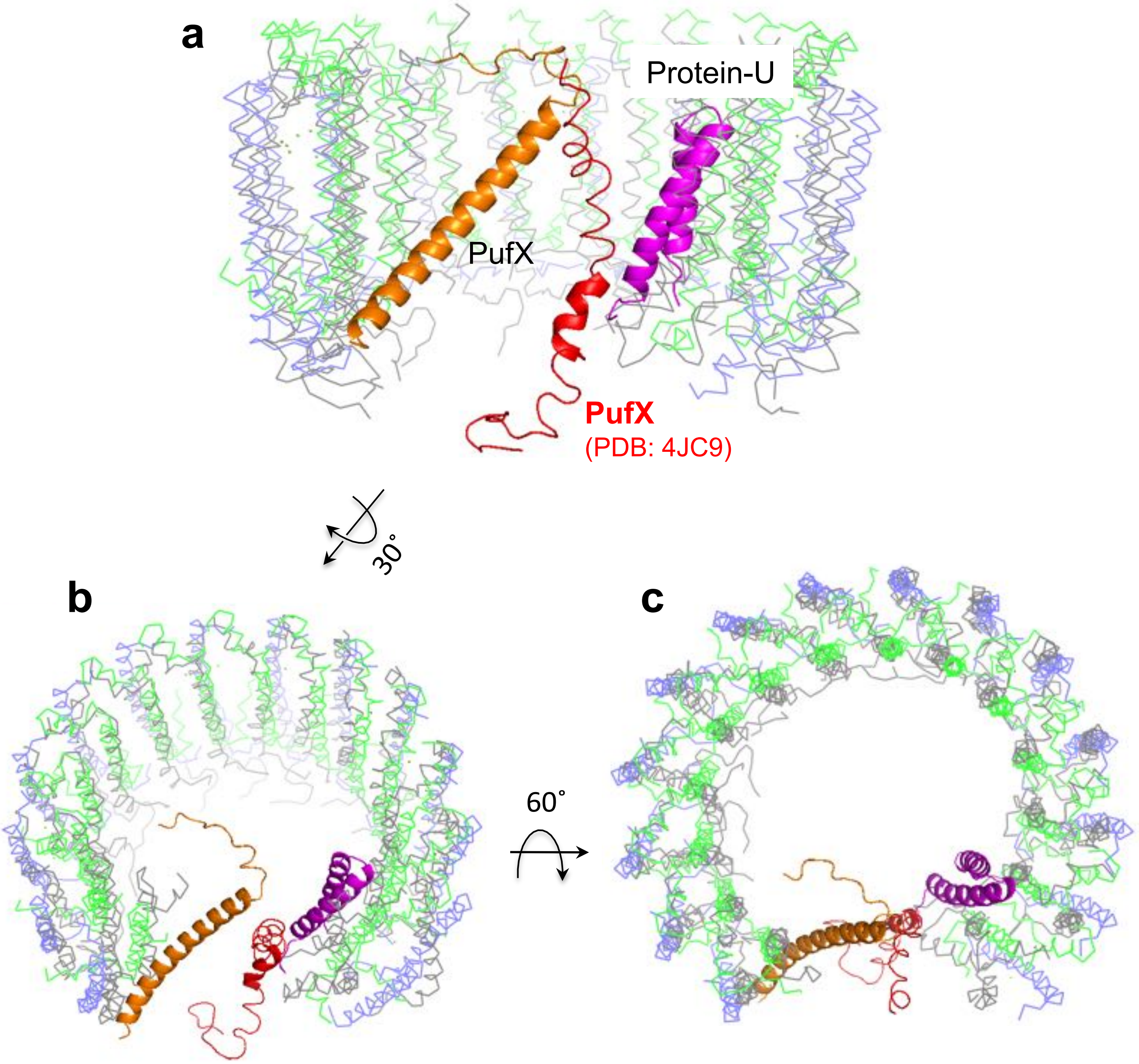

**Supplementary Fig. 5 Comparison of the PufX structure determined in this work (orange cartoon) with that previously reported in a dimeric LH1-RC structure from *Rba. sphaeroides* strain DBCΩG (red cartoon, PDB: 4JC9). (a)** Side view of superposition of the Cα carbons of the LH1 αβ-polypeptides between the structure in this work (LH1-α, green ribbon; LH1-β, slate-blue ribbon; Protein-U, magenta cartoon) and that in PDB 4JC9 (LH1-αβ, gray ribbons). **(b)** Tilted view of **(a)**. **(c)** Top view of **(a)** from the periplasmic side.

**a**

LH1-α (calculated Mw: 6837.11)
`formyl-MSKFYKIWMIFDPRRVFVAQGVFLFLLAVMIHLILLSTPSYNWLEISAAKYNRVAVAE`

LH1-β (calculated Mw: 5457.12)
`ADKSDLGYTGLTDEQAQELHSVYMSGLWLFSAVAIVAHLAVYIWRPWF`

PufX (calculated Mw: 7459.58)
`ADKTIFNDHLNTNPKTNLRLWVAFQMMKGAGWAGGVFFGTLLLIGFFRVVGRMLPIDENPAPAPNITG`

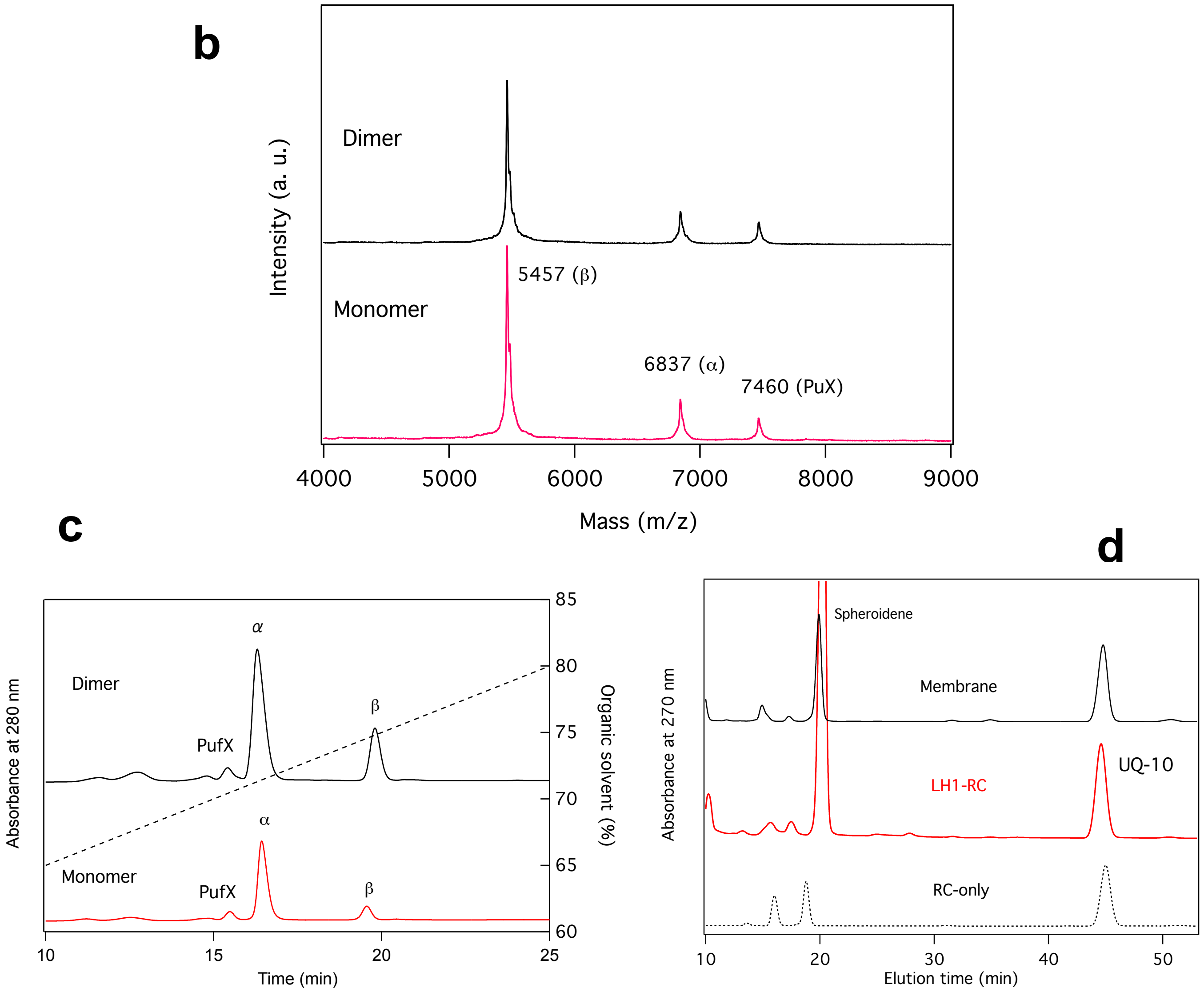

**Supplementary Fig. 6 Characterizations of the *Rba. sphaeroides* IL106 LH1-RC complex. (a)** Sequences of the expressed LH1-αβ and PufX polypeptides. **(b)** MALDI/TOF-MS spectra of dimeric and monomeric LH1-RCs obtained under the same conditions as described in Ref. 2. **(c)** Reverse-phase HPLC chromatograms (TSKgel, SuperODS, 4.6×100 mm, TOSOH) of the dimeric and monomeric LH1-RCs eluted at 25 ˚C by a gradient of 60–90% organic solvent consisted of acetonitrile/2-propanol (2:1) containing 0.1% trifluoroacetic acid. **(d)** Reverse-phase HPLC chromatograms (TOSOH, TSKgel ODS-80Ts, 4.6×250 mm) of the quinones and pigments from the membranes, purified LH1-RC and RC-only complexes isocratically eluted at 25 ˚C by 7:3 methanol/isopropanol at flow rate of 0.7 mL/min.

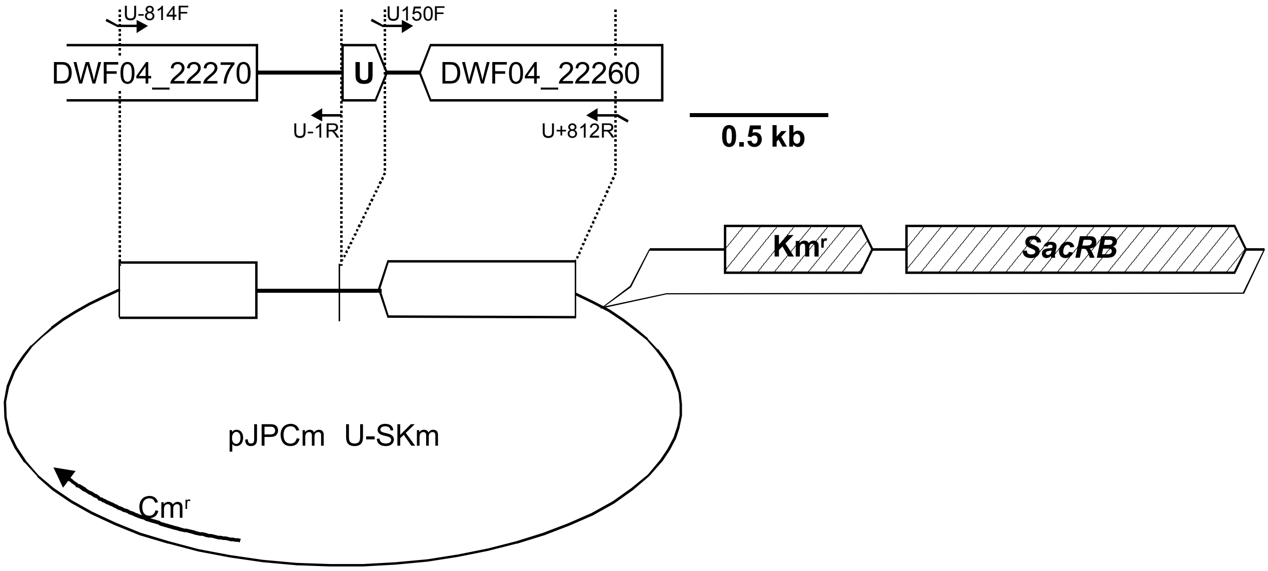

U-814F: 5'-TTGCATGCCTGCAGGTCCACCGCCACGAAGAGGAGGTTG
U-1R: 5'-GGTGCCTCCTTCAGATGCAAGC
U150F: 5'-TCTGAAGGAGGCACCGAACAGCAACTGACGGCACAGC
U+812R: 5'-GGGGATCCTCTAGAGTCGAGTTCACCGACTTCTTCGGCAAC

**Supplementary Fig. 7 Schematic representation of gene manipulation.** Genes are designated by open boxes with arrow heads showing the direction of transcriptions. The gene encoding the protein-U is labeled by "U", which is flanked by two ORFs (DWF04_22260 and DWF04_22270 by reference to NCBI database; QRBG01000031) on the complementally strand. Small arrows represent oligonucleotide primers used for PCR. Tabs at the ends of these arrows show additional sequences used for the ligation to the specified DNA fragments. Nucleotide sequences of these PCR primers are shown on the bottom, in which the part of the ligation-tab is underlined.

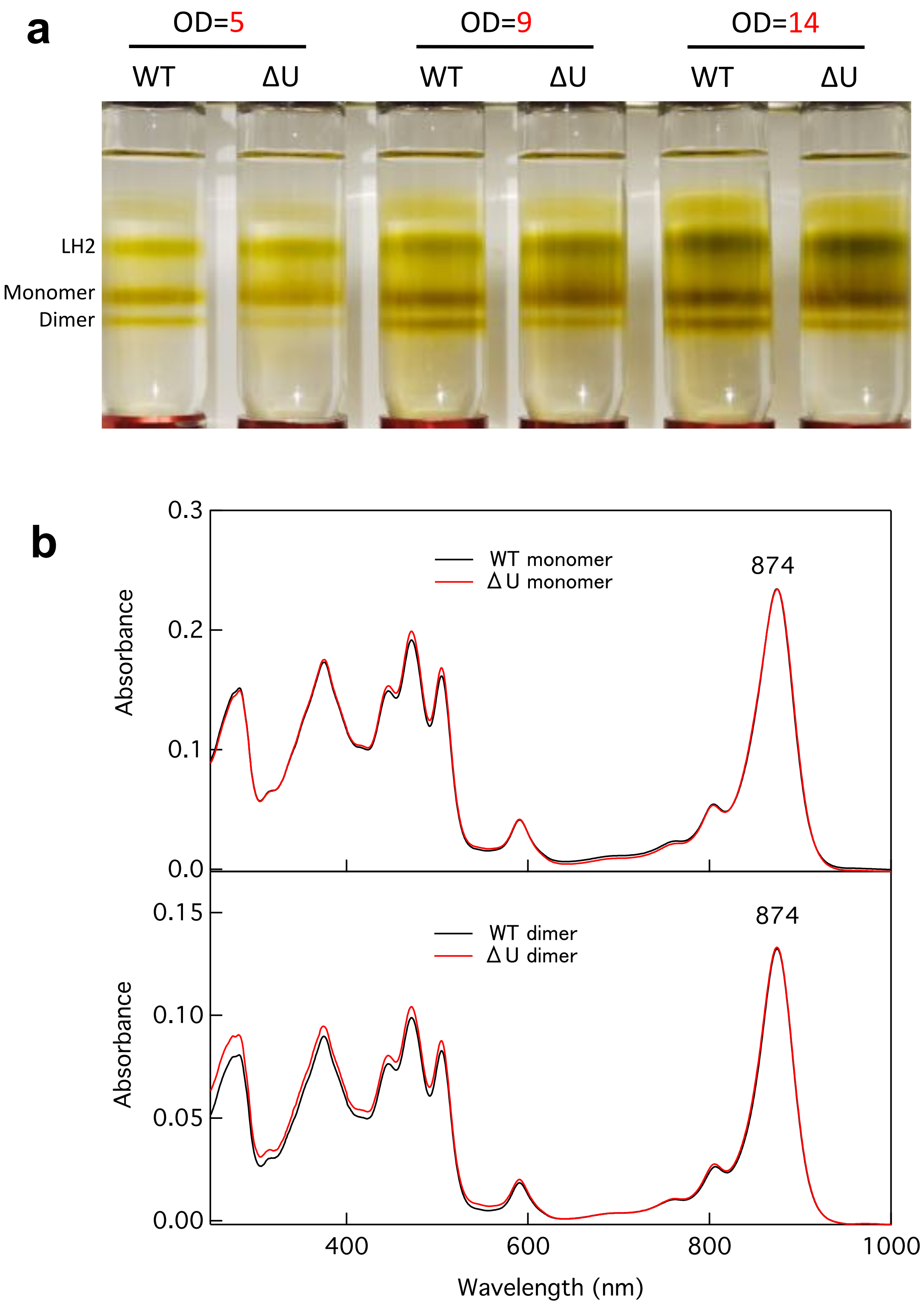

**Supplementary Fig. 8 Characterizations of the protein-U-deleted *Rba. sphaeroides* strain IL106ΔU. (a)** Sucrose density gradient (10–40% w/v) centrifugations of the solubilized pigment-protein complexes from wild-type (WT) and protein-U-deleted (ΔU) *Rba. sphaeroides* IL106 membranes. 1mL of the solubilized solutions with the concentrations (OD at 850 nm) indicated was loaded on the sucrose solution in each tube. **(b)** Normalized absorption spectra of the monomeric (*upper*) and dimeric (*lower*) WT and ΔU LH1-RCs collected from the sucrose density gradient solutions.

**a**

| Species name | Protein-U | PufX | Oligomeric state | References |
|---|---|---|---|---|
| ***Rba. sphaeroides* f. sp. *denitrificans* (IL106)** | **WP_002721225** | **WP_069333028** | **Monomer/Dimer** | *EMBOJ* 1999 18:534 |
| *Rba. sphaeroides* 2.4.1. (NCIB 8253) | WP_002721225 | WP_002720419 | Monomer/Dimer | *Nature* 2004 430:1058 |
| *Rba. sphaeroides* ATCC 17025 | WP_085996593 | WP_011909039 | | |
| *Rba. sphaeroides* ATCC 17029 | WP_002721225 | WP_002720419 | | |
| *Rba. sphaeroides* KD131 | WP_002721225 | WP_002720419 | | |
| *Rba. sphaeroides* WS8N | WP_002721225 | WP_002720419 | | |
| *Rba. azotoformans* | WP_085996593 | WP_011909039 | Monomer/Dimer | *BBA* 2012 1817:336 |
| *Rba. johrii* | WP_002721225 | WP_069333028 | | |
| *Rba. megalophilus* | WP_002721225 | WP_002720419 | | |
| *Rba. ovatus* | WP_176504535 | WP_097030925 | | |
| *Rba. sediminicola* | WP_085996593 | WP_145104437 | | |
| *Rba. blasticus* DSM 2131 | | WP_181318217 | Monomer/Dimer | *JBC* 2005 280:1426 |
| *Rba. capsulatus* B6 | | WP_013066439 | Monomer | *BBA* 2012 1817:336 |
| *Rba. veldkampii* DSM 11550 | | WP_107324827 | Monomer | *Struct*. 2007 15:1674 |
| *Rba. vinaykumarii* | | WP_076363364 | Monomer | *BBA* 2012 1817:336 |
| *Rba. aestuarii* | | WP_076484814 | | |
| *Rba. flagellatus* | | WP_149588102 | | |
| *Rba. maris* | | WP_097069514 | | |
| *Rba. thermarum* | | WP_128514093 | | |
| *Rba. viridis* | | WP_110805120 | | |
| *Cereibacter changlensis (Rba. changlensis)* | | WP_107664449<br>WP_136793197 | Monomer/Dimer | *BBA* 2012 1817:336 |
| *Rhodobaca bogoriensis* LBB1 | | WP_071479740 | Monomer/Dimer/Trimer | *Phil. Trans. R. Sci.* 2012 367:3412 |

**b**

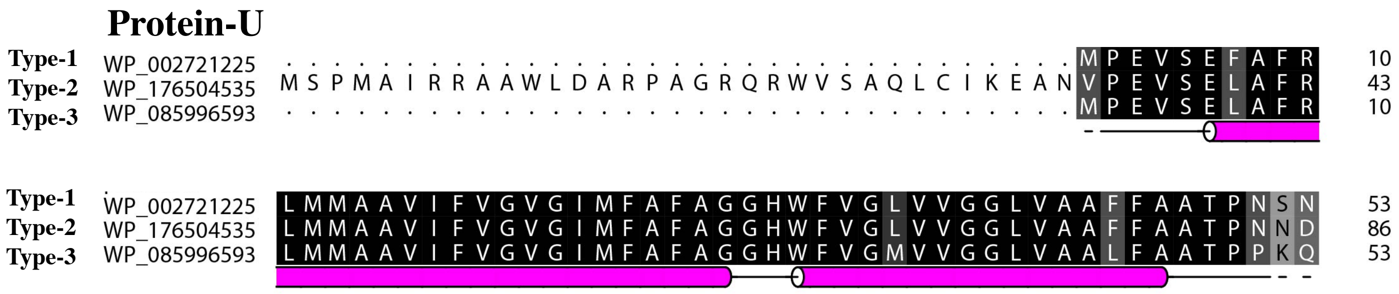

**Supplementary Fig. 9 Distributions of the protein-U and PufX in the genus *Rhodobacter*. (a)** Distributions of the protein-U and PufX (indicated by Protein ID) in the genome database of genus *Rhodobacter*. **(b)** Sequence comparison of the protein-U between the *Rba. sphaeroides* IL106 (WP_002721225) and others using the CLUSTAL X. The background of residues are colored in gray scale by similarity (Black: identical, White: non-conserved).

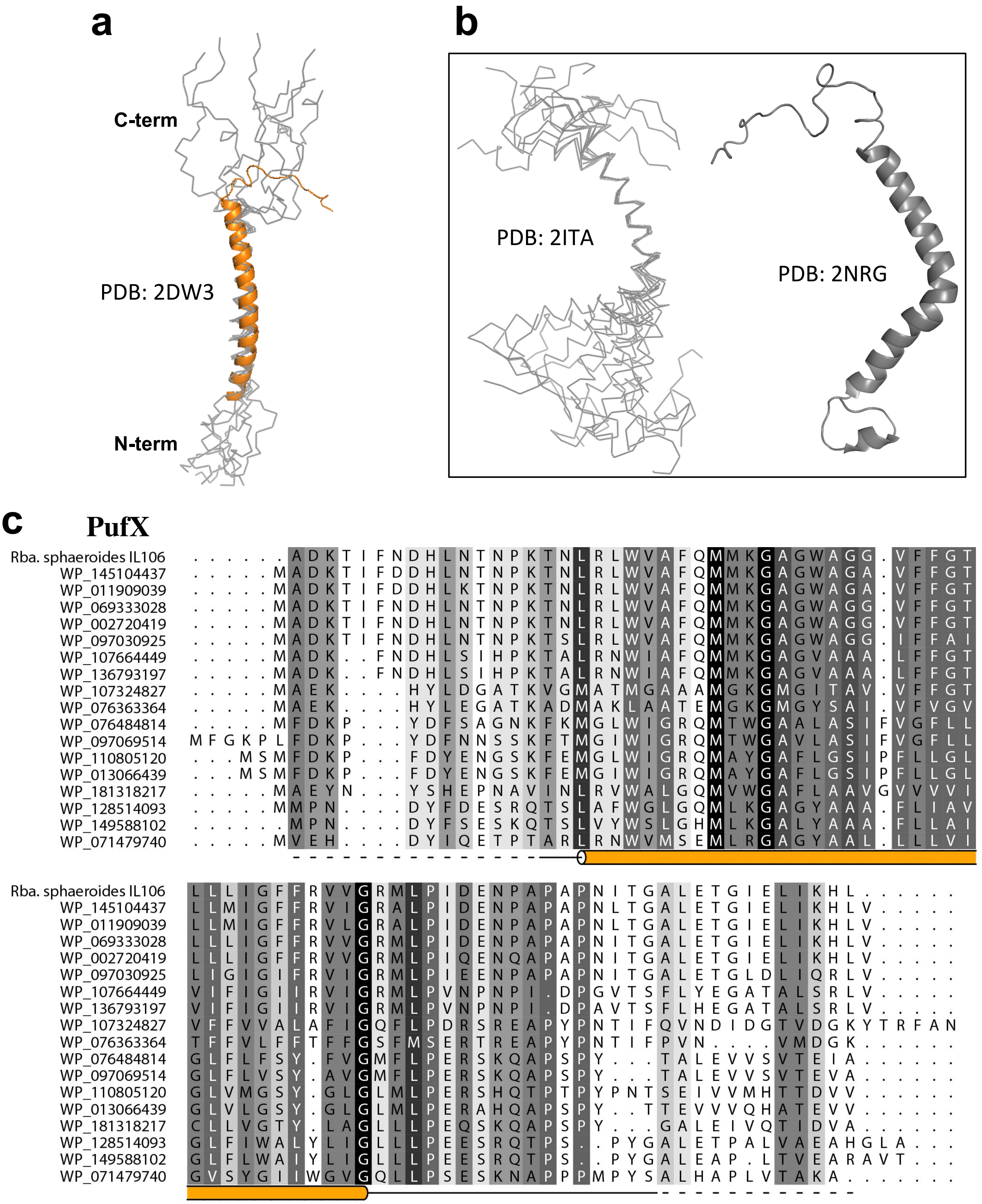

**Supplementary Fig. 10 Comparisons of the PufX structures and sequences. (a)** Superposition of the transmembrane domains for the *Rba. sphaeroides* PufX structures determined by cryo-EM (this work, colored) and solution NMR (gray ensemble, PDB: 2DW3). **(b)** Alternative solution NMR structure (PDB: 2ITA for the ensemble; PDB: 2NRG for the minimized average) of the *Rba. sphaeroides* PufX polypeptide. **(c)** Sequence alignment of the PufX and PufX-like polypeptides using the CLUSTAL X. The background of residues are colored in gray scale by similarity (Black: identical, White: non-conserved).
